# Seasonal variation of fermentation rate in *Saccharomyces spp*. (Ascomycota)?

**DOI:** 10.1101/285312

**Authors:** Dagmar Tiefenbrunner, Helmut Gangl, Ksenija Lopandic, Wolfgang Tiefenbrunner

**Author notes:** Correspondence should be addressed to W.T.

## Abstract

Yeast species of the genus *Saccharomyces* show some reaction to visible light – although they lack photo pigments and the typical clock genes of fungi – that can be explained by damage of the cytochrome electron transport chain of the mitochondria. Evidence for a circadian clock, entrainable by cyclic environmental stimuli, exists for periodic changing temperature and light as zeitgeber. Whether seasonality follows from the existence of a circadian clock – which is a necessary requirement for annual rhythms – remains unknown.

Due to an accidental observation, we were able to show that fermentation taking place in complete darkness and at constant temperature is influenced in some yeast strains by the history of the inoculum culture. Using yeast cultures growing on agar plates and exposed to diffuse daylight for three weeks either in March or in May as inoculum, leads to significantly different fermentation rates in the inoculated grape juice in both months: rates are higher in March when day length is shorter than in May. In must inoculated with cultures that grew in darkness or daylight, respectively, higher fermentation rates occur by the former. Other yeast strains react to artificial white light in the same way.

We used strains of *S. cerevisiae, S. eubayanus, S. kudriavzevii, S. uvarum* and furthermore hybrid strains of two or even three of these species. The most pronounced reaction to daylight was shown by the *S. eubayanus* x *S. uvarum* x *S. cerevisiae* hybrid, followed by *S. cerevisiae* x *S. kudriavzevii* hybrids, *S. eubayanus* and *S. cerevisiae. S. uvarum* was sensitive to artificial white light.

These observations can hardly be explained by some kind of photo damage because they base on an effect that persists through many cell division cycles after yeasts were exposed to light. If it really represents seasonality epigenetic memory is likely involved, since fermentation lasts for many days and yeast generations. If the existence of a circadian clock and seasonal behaviour in *Saccharomyces* is confirmed these yeasts could become an important tool in basic research concerning epigenetic memory.

## Introduction

In their review of photobiology of fungi, Roenneberg & Merrow 2001 stated that little is known about photoperiodism and seasonality in fungi in general and yeasts in special. This holds true by now, especially concerning yeasts, although some remarkable progress was made (e.g. Eelderink-Chen et al. 2010, Robertson et al. 2013).

Photoperiodic response needs light reception, but *Saccharomyces cerevisiae* (Ascomycota) cannot be seen as photoresponsive fungus because it lacks the photoreceptors and - pigments that many other organisms own, like rhodopsins, cryptochromes, phytochromes or white-collar 1. In many fungi, e.g. *Neurospora*, photo responses are controlled by two loci, white-collar-1 and white-collar-2 (Degli-Innocenti & Russo 1984, Russo 1988), the gene products of which are transcription factors. These clock genes are absent in *Saccharomyces* and hence it was doubted that baker’s yeast possesses a circadian clock like the one that regulates gene expression, metabolism and daily behaviour in other organisms. However, although *S. cerevisiae* has no orthologs of the transcription factors (clock genes) that mediate circadian regulation in fungi or animals and none of the photosensitive pigments that are necessary to entrain a circadian clock, it is well known that *S. cerevisiae* responds to visible light (e. g. Sulkowski et al. 1964, Ninnemann et al. 1970, Woodward et al. 1977, Edmunds et al. 1978, 1979a & b, Robertson et al. 2013).

Visible light has a significant impact on yeast metabolism, especially at low temperatures. Woodward et al. 1977 demonstrated that growth rate as well as sugar and amino acid transport were inhibited at 12°C and light intensities in excess of 1250 lx whereas there was no inhibition at 20°C. Even earlier, inhibition of protein synthesis by low-light intensities (Sulkowski et al. 1964) and respiration (Ninnemann et al. 1970) were reported. Ninnemann et al. (1970) recognized that oxidized cytochrome a_1_a_3_ is required for light inhibition. Thus it is likely that many of the effects of light on yeast cells can be explained by damage of the cytochrome electron transport chain and reduced mitochondrial activity. This assumption is confirmed by the fact that treating yeast with the electron transport inhibitor sodium azide has similar effects on yeast respiratory oscillation – as a sensitive measurement of metabolism – as visible light. Furthermore Robertson et al. 2013 showed that light can modulate cellular ultradian rhythmicity and that budding yeast can exhibit light-anticipatory behaviour that may be necessary in nature for protection against photo damage.

Evidence for the existence of a circadian clock that can be entrained by a cyclic environmental stimulus (zeitgeber) in *S. cerevisiae* were found by Eelderink-Chen et al. 2010 in experiments performed in a fermentor culture system that maintained cells in a nutritionally stable environment. The environmental stimulus was a periodically changing temperature so that lack of yeast photoreceptors didn’t matter. A circadian clock that can be entrained is the requirement for seasonal changes in behaviour or metabolism but as far as we know no hints for seasonality in yeasts were published up to now.

Here we report an accidental observation that is not a proof of but might be a clue to seasonal behaviour in *Saccharomyces* species. Surprisingly the behavioural changes are light induced but are independent of respiration. During four years we observed fermentation of grape juice in March and May and found that fermentation rate was always higher in March. These yeast strain dependent differences were especially amazing since fermentation took place in complete darkness and at constant temperature, only the cultures on agar plates used for inoculating were exposed to daylight. Thus epigenetic memory is likely involved into this phenomenon (e. g. Jaenisch & Bird 2003).

## Materials and Methods

### Yeasts

Yeast samples were taken out of liquid nitrogen three weeks before start of the experiment and were transferred to GYP agar plates (2% glucose, 1% peptone, 0.5% yeast extract) where they remained at room temperature. The Petri dishes were exposed either to diffuse daylight, or to artificial “daylight” or were kept in darkness, respectively. For production of artificial “daylight” (artificial white light) a Phototherapy Unit TL90 WL10, (Beurer GmbH, Ulm, Germany), was used. The unit has a light area of 51 cm x 34 cm and a colour temperature of 6500 Kelvin with an intensity of 10000 LUX at a distance of 15 cm. Petri dish distance was 45-50 cm. Temperature near the agar plates was 2°C above room temperature.

In the first year (2014) manifold yeasts were used (Tab. 1): four strains of *S. cerevisiae*, one *S. eubayanus, S. kudriavzevii* and *S. uvarum*, one *S. cerevisiae* x *S. eubayanus* hybrid, six hybrid strains *S. cerevisiae* x *S. kudriavzevii*, six strains *S. cerevisiae* x *S. uvarum*, one *S. kudriavzevii* x *S. uvarum* hybrid and one hybrid of *S. eubayanus* x *S. uvarum x S. cerevisiae*. Later on a subset of these yeasts was utilized: three strains of *S. cerevisiae* (commercial strain ‘Pannonia’, HA2796, HA1869), two of them isolated from fresh or fermenting grape juice, one of *S. eubayanus* (HA2841), isolated from the fruiting body of *Cyttaria hariotii* and one of *S. uvarum* (HA2786), gained from wine must, Tokaj wine region, Hungary.

**Table 1:**
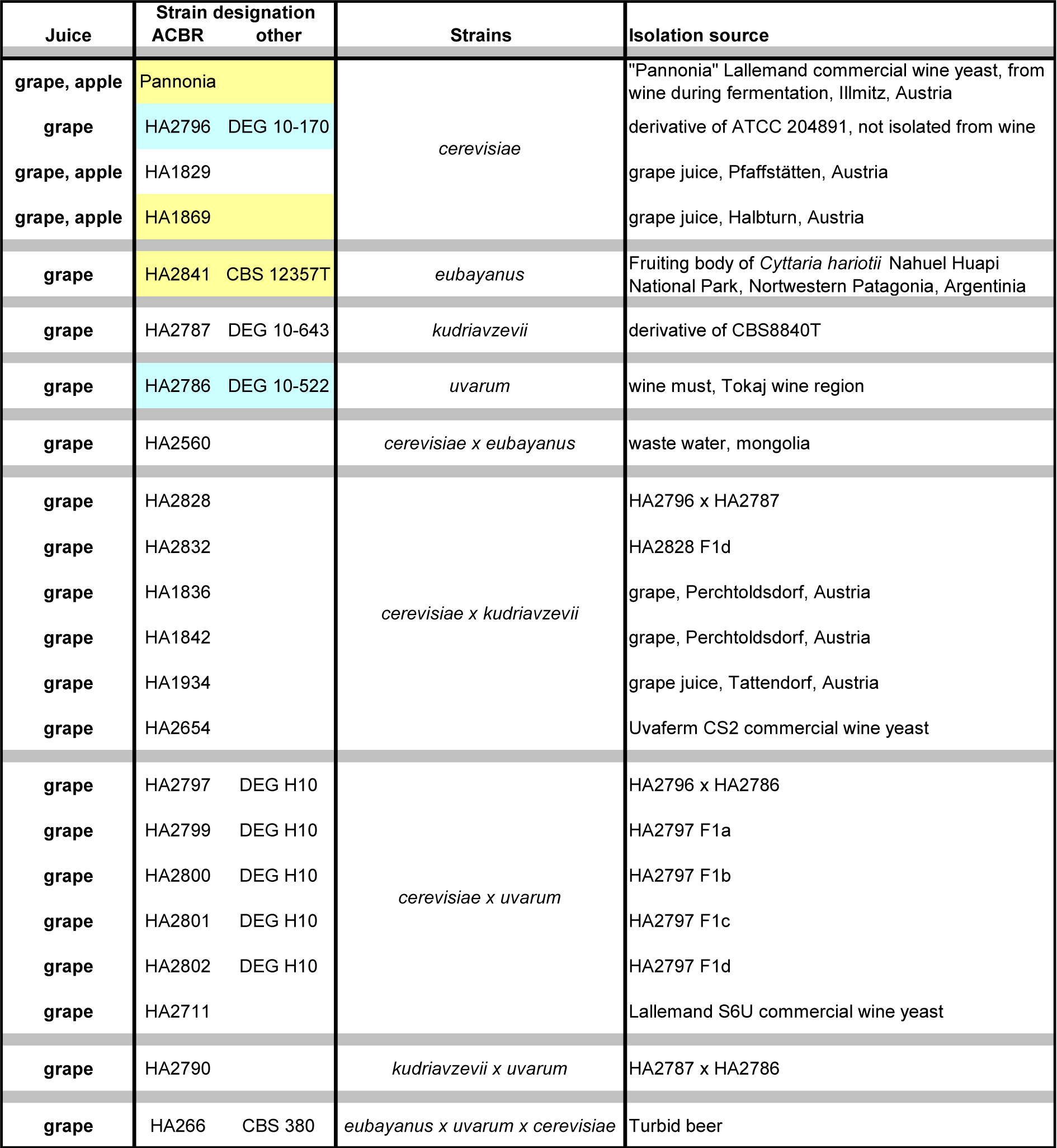
Yeast strains (Saccharomyces spp. and hybrids) used in the experiments. After 2014 only the highlighted strains were utilized (yellow: daylight sensititve strain, aqua: not sensitive to daylight). References: HA2796, HA2787, HA2786, HA2828, HA2832, HA2797, HA2799, HA2800, HA2801, HA2802, HA2654: Pfliegler et al. 2014; HA1829, HA1869, HA1836, HA1842: Lopandic et al. 2007; HA1934, HA2560: Stübiger et al. 2016; HA266: Nguyen et al. 2011; HA2841: Libkind et al. 2011; HA2711: Pérez-Torrado et al. 2015.

### Microvinification

Yeasts were inoculated in separate flasks. Microvinifications were carried out in March and May 2014-2017 in 300 ml Erlenmeyer flasks filled with 250 ml white grape juice (Grüner Veltliner) or apple juice in darkness at a standardised temperature of 20°C. Fermentation progress was monitored by determining the weight loss caused by the production of CO_2_. Transpiration loss was documented in a separate flask and considered. Fermentation lasted for about one month before wine chemical analysis eventually started. The resulting wines were tested olfactory. Odour and taste of all wines were acceptable, indicating that no bacterial or other contamination had occurred.

### Determination of fermentation rate

To determine fermentation rate a nonlinear regression analysis for the weight loss due to CO_2_ production was performed. The data were fitted to the Verhulst equation since metabolic turnover should mainly follow the population growth of the yeast in the flask. Three parameters α, β, γ can be condensed out of the logistic function for each original data set: y(t) = γ/(1+β exp(αt)). The Verhulst equation describes the growth of a population when resources are limited; its course is sigmoid (Hofbauer and Sigmund 1984). The parameter α can be used to estimate the fermentation rate. The coefficient of determination (R^2^), describing how well the data fit to the estimated function, can be used to see whether the fermentation occurred harmoniously, or was disturbed or interrupted. Although in our experiments R^2^ indicates that fitting quality of the function to the data was generally very high (0.959≤ R^2^≤0.998), there were two systematic deviations (Fig. 1), one at the beginning and one at the end of the fermentation process. We argue that at the beginning there is enough oxygen in the grape juice that respiration is possible, which leads to a higher metabolism and population growth than can be explained by pure fermentation and thus by the Verhulst function. Later on the medium is oxygen deprived: respiration is not possible any more and fermentation takes place. At the end of the process, when no more sugar is available, fermentation stops. However, some metabolic processes remain since weight loss in this phase is higher than can be explained by transpiration loss. Because weight loss is now linear it is likely that respiration occurs, because oxygen diffusion is proportional to medium surface and although there is no more sugar available, other resources remain.

**Fig. 1:**
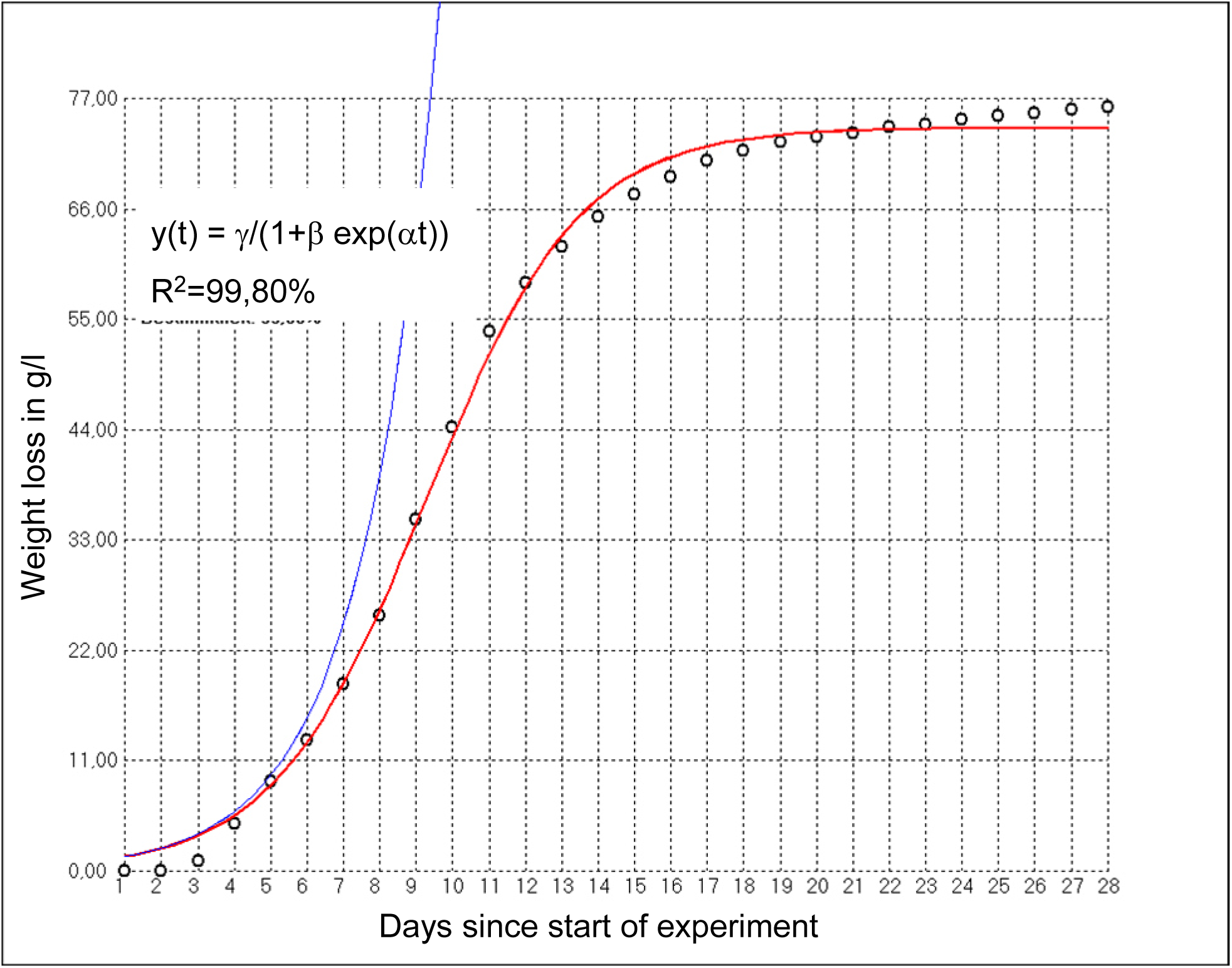
Fitting of the Verhulst function to the weight loss during fermentation due to CO_2_ production in one of the flasks. The red curve corresponds to the Verhulst function y(t) = γ/(1+β exp(αt)) (blue curve: y(t) =exp(αt)). Although the coefficient of determination shows a very high value, fitting at the beginning an end of the process is not perfect.

Since the middle part of the fermentation is most important for determining the rate, we ignored the systematic deviations. Data had to be prepared, because, for example, not the weight of the entity but the weight loss since the beginning of the experiment had to be taken into account. Transpiration loss and initial weight differences of the grape juices in the flasks had to be considered.

We utilized our own software for regression. For nonlinear regression, in many cases special algorithms for solving the non-linear least squares problem must be used since analytical solution is not possible, examples of which are the Gauss–Newton algorithm or the Levenberg–Marquardt algorithm. Here we utilized the function space concept (Fig. 2), since it doesn’t depend that much on initial conditions.

**Fig. 2:**
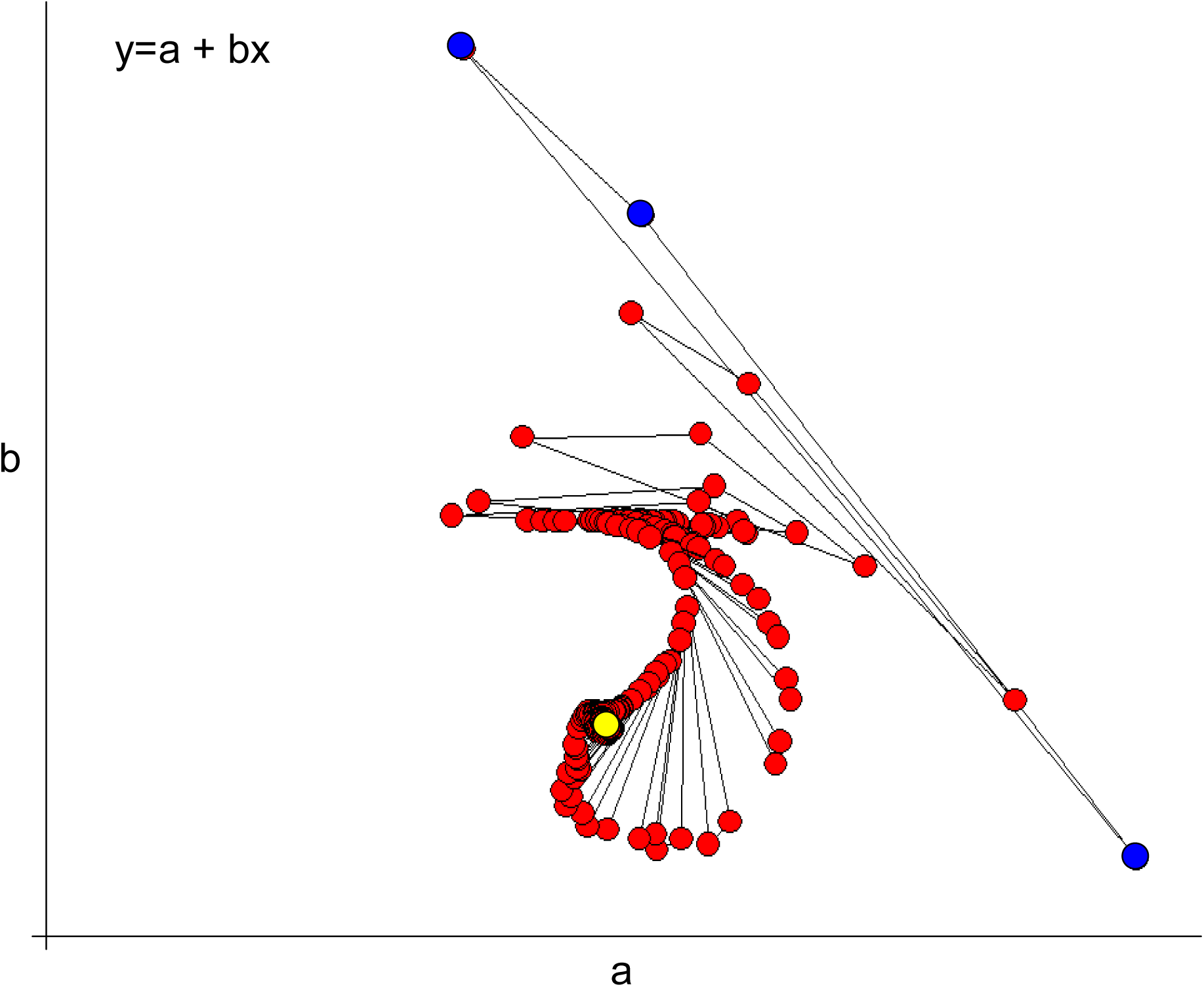
The amoeba search procedure for finding the absolute terms of a function (in this case a straight line or any function with two absolute terms a and b). Blue: randomly chosen initial “amoeba”. Yellow: target. Red: successive positions of the “amoeba” during the search procedure.

Any function can be seen as point in a multidimensional space where the number of dimensions is given by the quantity of absolute terms in the function. The values of the absolute terms give the position of the point.

To find any point with unknown position – and thus the values of the absolute terms (which resembles the regression problem) – one can use randomly set search points, the number of them must be at least one more than the quantity of dimensions. Calculating the sum of least squares for these points (blue in Fig. 2) gives the relative distance to the target point (yellow) or function for all of them. Using vector addition and the “distance” between the search points, the position of them is changed little by little (red in Fig. 2), until the target is reached. For movement (change of position) the information is used, which point is nearer to the target and how much it is nearer. Since this is presumably the way an amoeba finds to its resources the procedure is named after this protist.

### Additional statistics

In order to assess differences between wines produced at two terms (March and May) during the year by hybrid yeasts and their parental species, the fermentation rate data and the ones of the basic chemical analyses were exported to Statgraphics Centurion Version XV software (StatPoint Inc., Waarenton, VA, USA, 2005). Mean comparison procedures for paired samples (t-Test and Wilcoxon signed rank test) were performed as well as those for independent samples (t-Test, Wilcoxon-Mann-Whitney). After 2014 multiple replications of each experiment were performed in both months and hence a new situation for statistical analysis emerged, which shall be explained by an analogy. Suppose someone wants to find out whether human males and females differ in body height on average. The person can take independent samples (e.g. randomly chosen 100 adult women and men) or paired samples, e.g. 100 pairs of siblings. However, the statistician can also take more siblings, not just pairs, for comparison. Since we do not know any test for this kind of “group comparison”, we designed our own permutation test. Within the group (the family in our analogy; actually all fermenting replications with the same yeast from the same year and month belong to the same group) we changed the pairing randomly and did so one million times to get the probability that the differences between the fermentation rates of March and May were as high or higher than observed either for the original data or their ranks. This test can be used for paired and independent samples, too (paired: group contains only one member of each variant; independent: group contains all members of each variant).

Chemical profiles were evaluated using principal component analysis (Hartung and Elpelt 1999) using ViDaX Version IV (LMS-Data, Trofaiach, Austria, 2005).

## Results and Discussion

On 10th March 2014 we started an experiment with the goal to find out which interspecies yeast hybrids of the genus *Saccharomyces* are qualified for wine production under several conditions (Lopandic et al. 2016, Gangl et al. 2017). Due to technical problems we had to repeat the fermentation process which happened almost exactly two months later. Although not necessary for the experiment, with some delay one of us calculated the fermentation rates of the failed March-experiment and compared it with the ones of May. At the first glimpse the result was not delightful since correlation of March and May fermentation rates was low (R=0.46) which was problematic for the significance of the question concerning the quality of the yeasts for wine production. However, at the second glimpse something interesting appeared: in most cases fermentation rates in March were higher than those of May (Fig. 3 and Fig. 4a), although the used grape (Grüner Veltliner 2013, 13,5° KMW) or apple juice (harvest 2013) was the same as well as the general conditions: fermentation took place at 20°C in complete darkness. This result is highly significant, the null hypothesis of the t-test for paired samples (26 pairs) that the mean March fermentation minus May fermentation equals 0,0 has to be rejected (P=0,0002, data do not deviate significantly from normal distribution), the nonparametric signed rank test gives qualitatively the same result (P=0,0003).

**Fig. 3:**
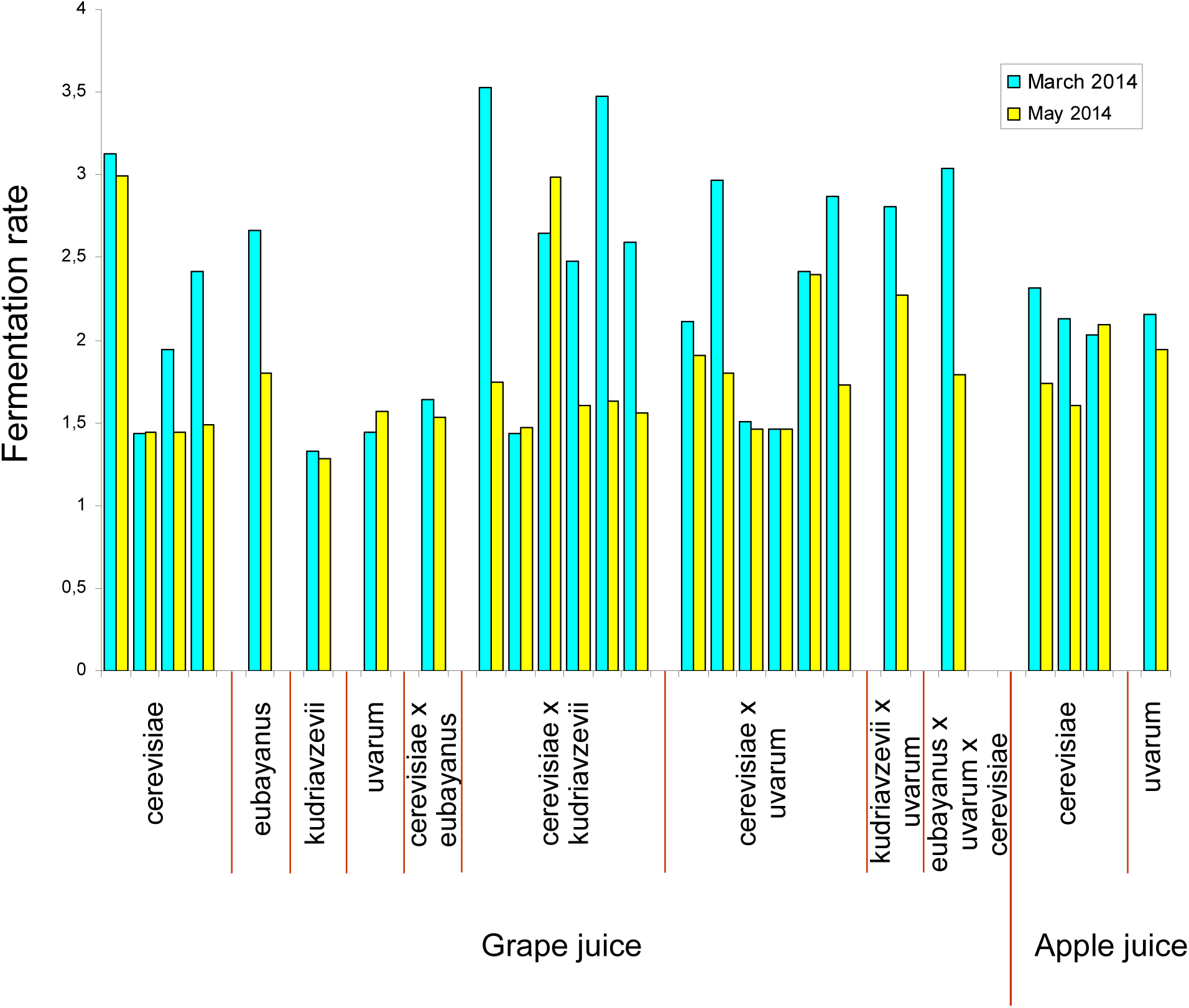
Fermentation rates of yeast strains. Fermentation took place in March and in May under the same conditions.

**Fig. 4:**
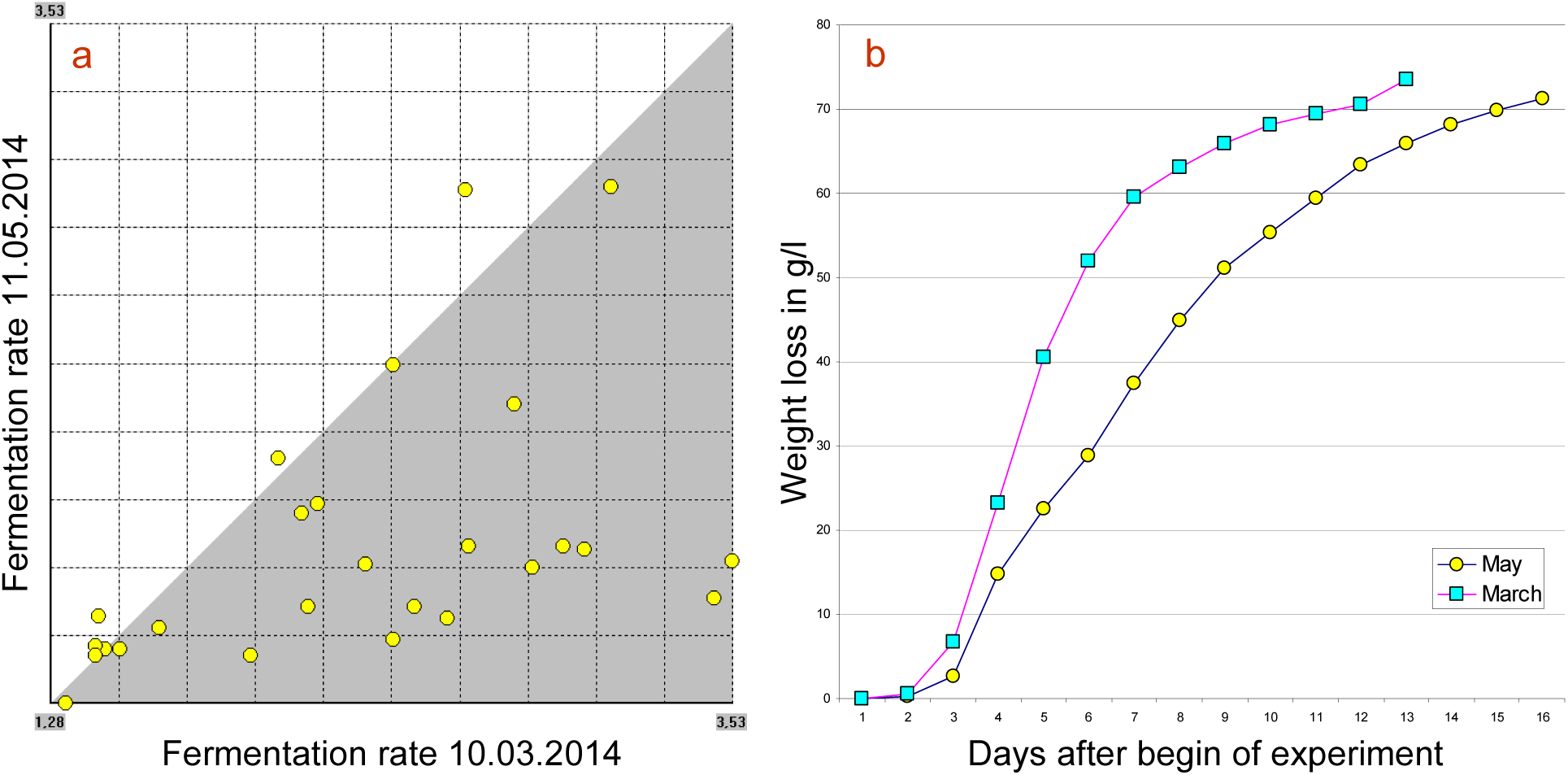
**Fig. 4a:** Fermentation rates with fermentation begin at 10.03.2014 presented against the ones with starting time 11.05.2014; shaded area: March>May. **Fig. 4b:** Weight loss during S. eubayanus fermentation of grape juice (March: fermentation rate 2,66 and May: fermentation rate 1,8).

The disparity between March and May fermentation was not the same for all yeast strains. March to May relation was highest on *S. eubayanus* x *S. uvarum* x *S. cerevisiae* hybrid (1.7), followed by *S. cerevisiae* x *S. kudriavzevii* hybrids (1.54 on average) and *S. eubayanus* (1.48, Fig. 4b) in grape juice. In *S. cerevisiae* the relation was 1.25 in grape and 1.21 in apple juice in mean. HA1869 isolated in Halbturn had a more pronounced disparity than the other strains of the species in fermenting grape juice. In *S. kudriavzevii* x *S. uvarum* the relation was 1.24 as well as in *S. cerevisiae* x *S. uvarum* on average. The fermentation rate of *S. uvarum* in March was lower than the one in May, so the relation is only 0.97.

Although the observed difference was highly significant, it seemed necessary to probe repeatability in an experiment of smaller extend. We choose two yeasts that showed high seasonal differences of fermentation rates (Pannonia *S. cerevisiae* and HA2841 *S. eubayanus*) and two with low differences (HA2796, *S. cerevisiae*, the strain not isolated from wine or juice and HA2786 *S. uvarum*) and repeated the experiment during 2015 - 2017. In the last year we added one more yeast, HA1869, with pronounced disparity. 2014 to 2016 we used the same grape juice (harvest 2013) but had to change the fermenting medium in 2017 because nothing was left (white grape juice Grüner Veltliner, harvest 2016, 14,5°KMW). Apple juice fermentation was not repeated. With exception of 2014, for each yeast strain we performed two (2015, 2016) to three (2017) replications a year (Fig. 5 and Fig. 6).

**Fig. 5:**
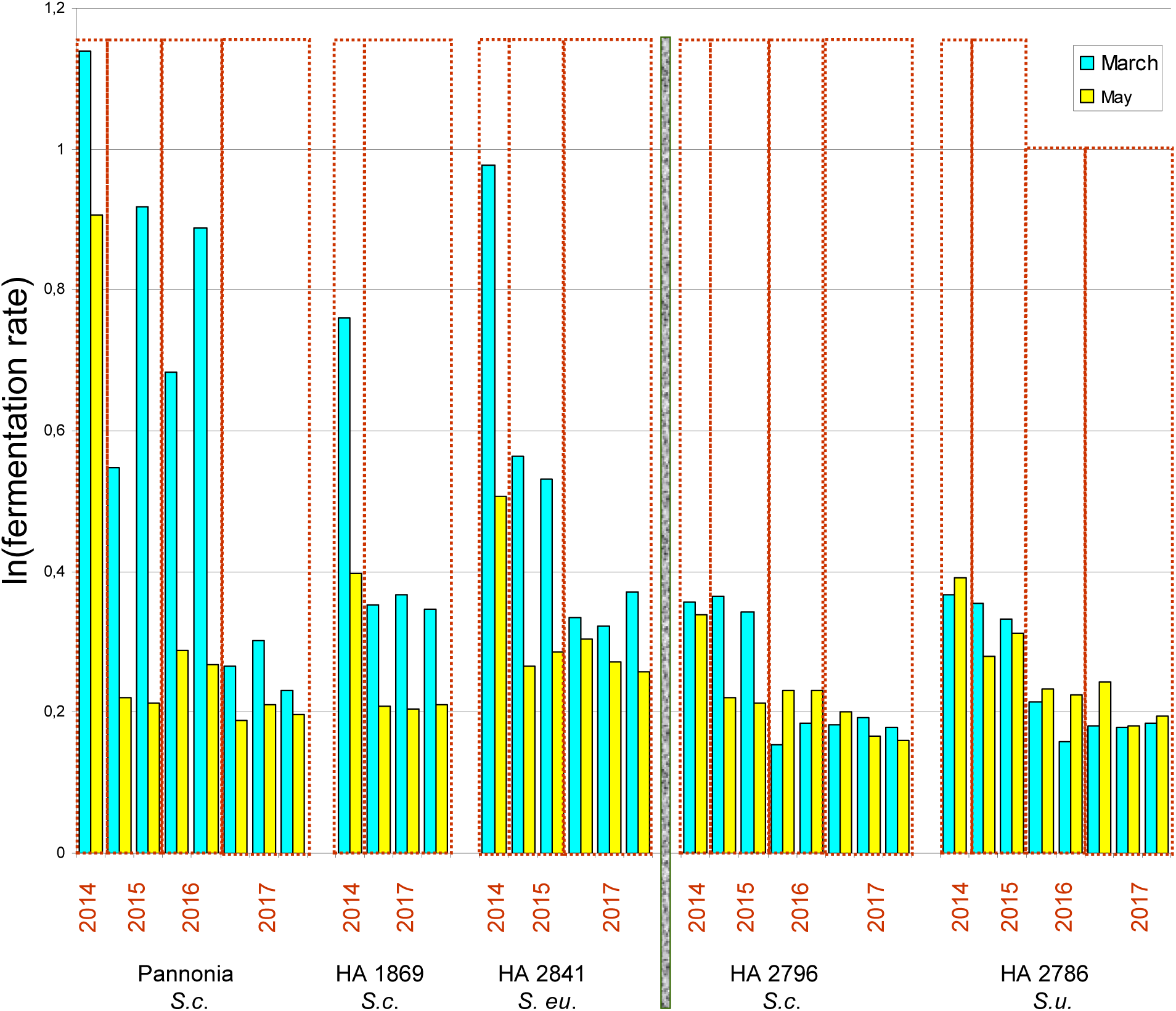
Comparison of fermentation rates in March and May during 2014 to 2017. The results for five yeasts are shown, three of them (left) sensitive to the date of the start of the experiment, two (right) not.

**Fig. 6:**
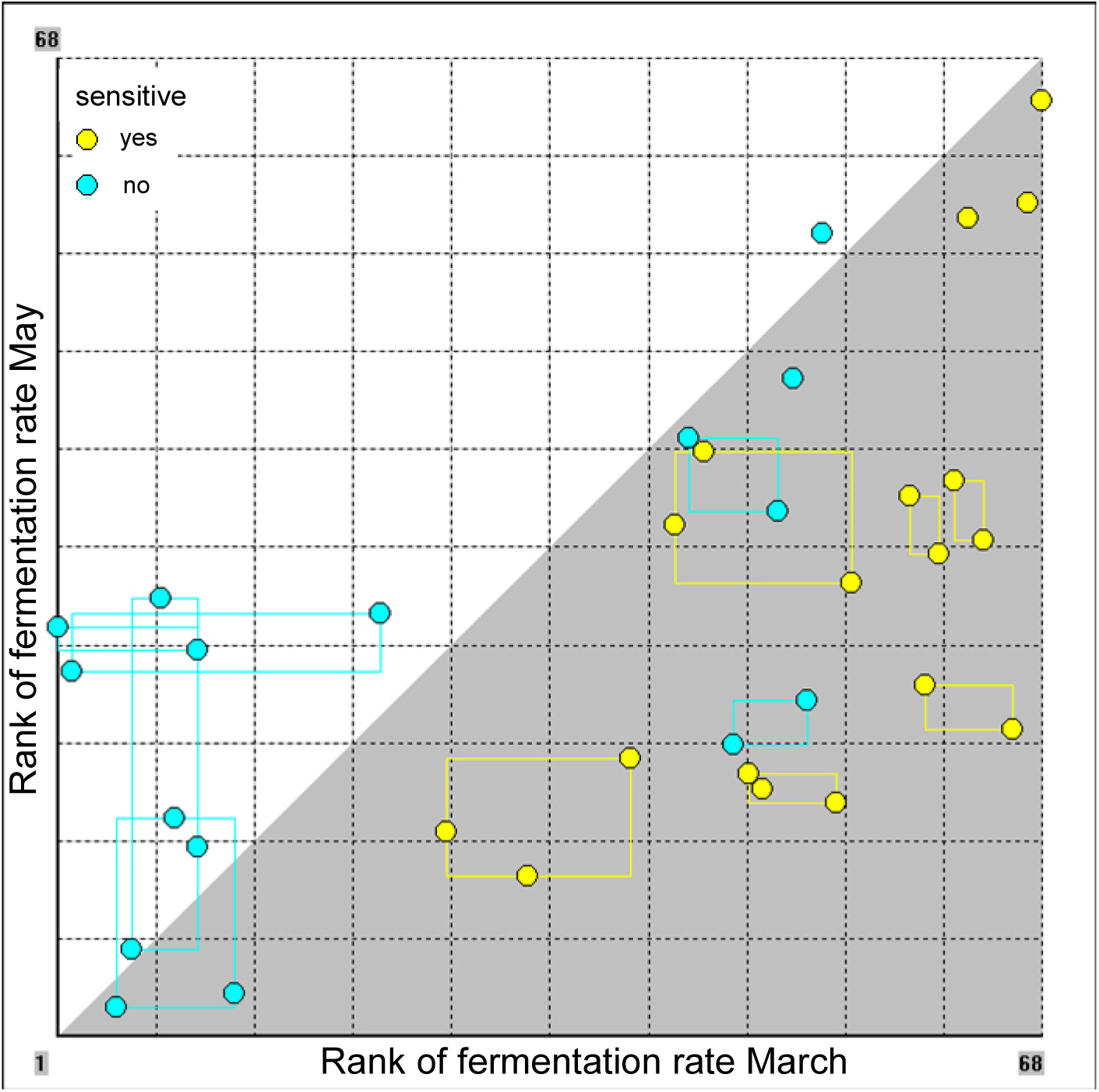
Comparison of the fermentation rates of March and May of several years (2014-2017). Ranks of values are used because of skewness of distribution. Sensitive yeasts always had higher fermentation rates in March. Groups (all fermenting replications with the same yeast from the same year) are marked by a rectangle. Shaded area: March>May. See also Fig. 5.

During four years none of the “sensitive” yeast strains (Pannonia *S.c*., HA1869 *S.c*. and HA2841 *S. eu*.) had a higher or equally high fermentation rate in May than in March (n=18 “pairs”) and hence in comparing the replications of each year not only the averages but all rates of March were higher than all of May from the same year. This is the reason that statistical tests denote a highly significant difference between fermentation in March and May, no matter whether the data are treated as grouped, paired or independent and whether original data or ranks are used (Tab. 2), although fermentation occurred slower in 2017, maybe because another grape juice had to be utilized. A generally lower rate could have some influence if data are treated as independent. With grouped (all replications with the same yeast fermented in the same year and month belong to one group) data – which is the best way to use all available information – P=0,00001 (permutation test).

**Table 2:** P-values of different statistical tests concerning the question whether there is a significant difference in fermentation rates of March and May in sensitive yeasts

| sensitive yeasts 2014 - 2017 |  |  |
| --- | --- | --- |
| permutation test | original data | ranks of data |
| grouped | 0.00001 | 0.00001 |
| paired | 0.00001 | 0.00002 |
| independent | 0.0067 | 0.00085 |
|  | t-test | signed rank test |
| paired | 0.00006 | 0.00021 |
|  | t-test | Wilcoxon-Mann-Whitney |
| independent | 0.00336 | 0.00029 |

In the yeast strains that are classified as not sensitive (HA2796 and HA2786) there is no significant difference between March and May fermentation (n=16), no matter which test was used (permutation test for grouped data P=0.7 for original data and P=0.48 for ranks). Thus yeast strains stay either sensitive or not through years.

Since the general conditions of the experiment were the same in March and May one wonders which influence caused the different behaviour. Before the beginning of fermentation, yeast cultures, provided for inoculation, grew on agar plates and were exposed to diffuse daylight; so day length could be perceived by the yeasts. To test this hypothesis we exposed a part of the yeast cultures for three weeks to darkness or artificial daylight (white light), respectively. In March the artificial length of the day we choose was that of 15^th^ of May in Vienna (15h 14min) and in May that of 15^th^ of March (11h 54min). Results are shown in Tab. 3 for sensitive yeasts and Tab. 4 for insensitive ones.

**Tab. 3:**
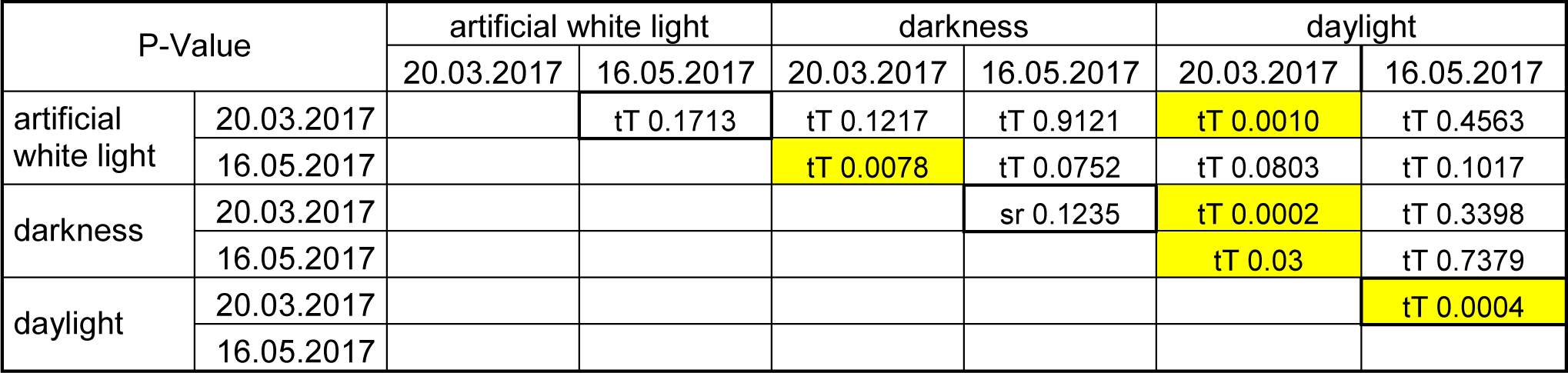
P-values for parametric and nonparametric tests for sensitive yeasts (Pannonia, HA1869, HA2841). Compared are fermentation rates. If the test requirement (normal distribution of data) is fulfilled the results of the t-test are shown, otherwise the ones of the signed rank test. tT: t-test for paired samples. sr: Wilcoxon signed rank test.

| P-Value |  | artificial white light |  | darkness |  | daylight |  |
| --- | --- | --- | --- | --- | --- | --- | --- |
|  |  | 20.03.2017 | 16.05.2017 | 20.03.2017 | 16.05.2017 | 20.03.2017 | 16.05.2017 |
| artificial white light | 20.03.2017 |  | tT 0.1713 | tT 0.1217 | tT 0.9121 | tT 0.0010 | tT 0.4563 |
|  | 16.05.2017 |  |  | tT 0.0078 | tT 0.0752 | tT 0.0803 | tT 0.1017 |
| darkness | 20.03.2017 |  |  |  | sr 0.1235 | tT 0.0002 | tT 0.3398 |
|  | 16.05.2017 |  |  |  |  | tT 0.03 | tT 0.7379 |
| daylight | 20.03.2017 |  |  |  |  |  | tT 0.0004 |
|  | 16.05.2017 |  |  |  |  |  |  |

**Tab. 4:** P-values for parametric and nonparametric tests for yeasts classified as not sensitive (HA2796, HA2786). Compared are fermentation rates. If the test requirement (normal distribution of data) is fulfilled the results of the t-test are shown, otherwise the ones of the signed rank test. tT: t-test for paired samples. sr: Wilcoxon signed rank test.

| P-Value |  | artificial white light |  | darkness |  | daylight |  |
| --- | --- | --- | --- | --- | --- | --- | --- |
|  |  | 20.03.2017 | 16.05.2017 | 20.03.2017 | 16.05.2017 | 20.03.2017 | 16.05.2017 |
| artificial white light | 20.03.2017 |  | sr 0.0360 | tT 0.0403 | tT 0.2445 | tT 0.2743 | tT 0.1590 |
|  | 16.05.2017 |  |  | sr 0.0360 | sr 0.0355 | sr 0.2945 | sr 0.0592 |
| darkness | 20.03.2017 |  |  |  | tT 0.2850 | tT 0.9677 | tT 0.1483 |
|  | 16.05.2017 |  |  |  |  | tT 0.6456 | tT 0.8675 |
| daylight | 20.03.2017 |  |  |  |  |  | tT 0.5587 |
|  | 16.05.2017 |  |  |  |  |  |  |

In 2017, the sensitive yeasts that were exposed to daylight before fermentation started (n=9 pairs) had always higher fermentation rates in March (average 1.38) than in May (average 1.26). This difference is significant (t-Test for paired data P=0,0004 Tab. 3; permutation test for grouped data P=0,004). As expected, no difference concerning daylight length could be seen in yeasts classified as not sensitive (n=6 pairs; t-test for paired data P=0.56; permutation test for grouped data P=0.65). Rates of fermentation were about 1.2.

There is no significant difference between fermentation rates of March and May concerning sensitive yeasts exposed to darkness or artificial white light, respectively, before fermentation started. Thus it was not possible to provoke different fermentation speed by using artificial light. Surprisingly, yeast classified as not sensitive showed differences in fermentation rates in the two months. Fermentation occurred faster (mean rate: 0.25) in May where artificial light exposure was shorter than in March experiment (mean rate: 0.17) and hence followed the trend that can be seen by sensitive yeasts and daylight. Significance was moderate (signed rank test P=0.036; P=0.031 permutation test for grouped data) although in all cases fermentation rates were higher in May (under the artificial day length of mid March), without exception. Thus it is likely that the yeasts classified as insensitive are not insensitive to light exposure in general but only to the diffuse day light we offered to them.

Sensitive yeasts treated darkness and artificial white light differently to daylight (Tab. 3), the fermentation was fastest if treated with darkness. “Insensitive” ones reacted differently only to artificial light and darkness (Tab. 3) but the rate differences were low.

The fermentation products of the March and May 2017 experiments were analyzed chemically. The wine parameters density, content of ethyl alcohol, fructose, glucose, titratable acid, volatile acids, tartaric acid, malic acid, citric acid, lactic acid and pH were determined and were used for a multivariate comparison of the daylight-variant (Fig. 7). Principal component analysis shows that PC1 distinguishes between March and May. Although there is no complete separation for all yeasts, it is for the sensitive ones. For each of them, March repetitions are further right (higher values of PC1). Concerning the insensitive ones, in HA2796 no separation along PC1 can be observed. Wines of March, when fermentation rates were higher, are much more similar then the ones of May as far as sensitive yeasts are concerned.

**Fig. 7:**
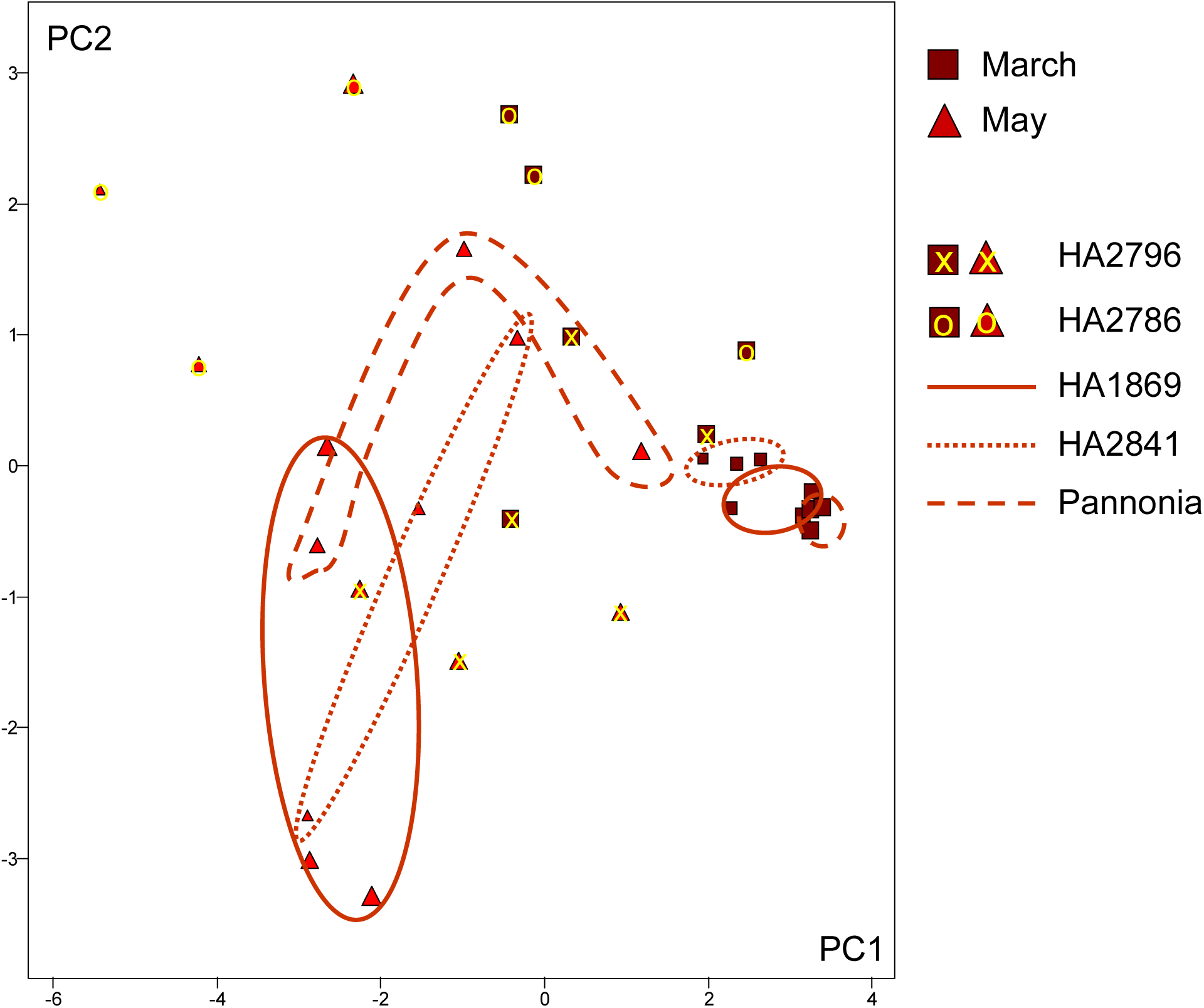
Three dimensional Principal Component Analysis of basic wine chemical parameters (PC3 can be seen through different size of symbols). Together PC1 to PC3 account for 85,79% of the variability in the original data.

The first principal component has the equation:

PC1=-0,34* Density + 0,31* Ethanol - 0,35* Fructose - 0,32* Glucose - 0,29* titratable acid + 0,28*pH + 0,31*volatile acids - 0,28* tartaric acid + 0,35* malic acid - 0,17* lactic acid - 0,28*citric acid

Thus, with the exception of lactic acid, all components contribute similar to PC1 value. March wines of sensitive yeasts have higher ethanol, volatile acids and malic acid content and a higher pH-value and lower density, sugar, titratable acid, tartaric acid and citric acid concentrations. Higher fermentation rates don’t necessarily presuppose higher alcohol concentrations since not only the speed but also the duration of fermentation is of importance but here we observed this relationship.

We have shown that yeast fermentation in grape juice in darkness and at constant temperature is influenced by length of daylight exposure to the inoculum culture in the weeks before fermentation starts. This strain dependent behaviour indicates that a circadian clock with light as zeitgeber exists which is the necessary requirement for seasonality. Although fermentation rates are higher if inoculum is held in complete darkness and shrinks with the length of daily light exposure the influence can hardly be a consequence of some damage because it remains through generations. Alternatively some epigenetic memory (heritable change in gene function that do not involve a change in the DNA sequence) is required to explain the observation. In this case the phenomenon could be of great interest for basic research if light dependent reaction can be initiated more successfully by artificial light that mimics daylight better. Whether artificial white light in our experiments induced the phenomenon in some yeast strains is not absolutely sure since temperature was also raised during culture growth on agar. In any case, for fermentation experiments in the lab, a detailed description of the history of the inoculum has to be made and as a standard it should be kept in complete darkness. Many questions remain: Which wavelengths of the light have the most pronounced influence and which intensity? How is the full year circle? What consequences have different temperatures during fermentation and the foregoing preparing part of the experiment? Is there any connection to the light effects on respiration? Why are there differences between yeast species and diverging strains of the same species?

In their review paper concerning seasonality and photoperiodism in fungi Roenneberg & Merrow 2001 start with a chronobiologist conversation in the famous Munich Schneiderbräu. They talk about how different the Bavarian beers are throughout the year (Starkbier in spring, Weißbier in summer, Wiesenbier in autumn and Weihnachstbier in winter) and wonder whether this could be because of yeasts being different at different times of the year. Although we cannot tell in this case, we can say by now that it might be. However, confirmation of yeast seasonality by a second lab that deals with chronobiology is urgently needed.

## Acknowledgements

The authors would like to thank Walter Pfliegler, Mathias Sipiczki and Ulziinyam Rentsendorj for providing strains. For discussion and support concerning method we thank Astrid and Martin Tiefenbrunner.

## References

Degli-Innocenti, F., Russo, V.E., Isolation of new white collar mutants of *Neurospora crassa* and studies on their behavior in the blue light-induced formation of protoperithecia, 1984, J. Bacteriol. 159, 757–761.

Edmunds Jr., L.N., Woodward J.R., Cirillo, V.P., 1978, Light effects in yeasts: Inhibition by visible light of growth and transport in *Saccharomyces cerevisiae* grown at low temperatures, J. Bacteriol 133, 692–698.

Edmunds Jr., L.N., Apter, R.I., Rosenthal, P.J., Shen, W-K, Woodward J.R., 1979a, Light effects in yeast: Persisting oscillations in cell division activity and amino acid transport in cultures of *Saccharomyces cerevisiae* entrained by light:dark cycles Photochem. Photobiol. 30: 595–601.

Edmunds Jr., L.N., Ulaszewski S., Mamouneas T., Shen, W-K, Rosenthal, P.J., Woodward J.R., Cirillo, V.P., 1979b, Light effects in yeasts: Evidence for participation of cytochromes in photoinhibition of growth and transport in Saccharomyces cerevsiae cultured at low temperatures, J. Bacteriol. 138, 523–529.

Edmunds Jr., L.N., 1980, Blue-Light Photoreception in the Inhibition and Synchronization of Growth ans Transport in the Yeast *Saccharomyces*, In: The Blue Light Syndrome (Ed. H. Senger), Springer.

Eelderink-Chen, Z., Mazzotta, G., Sturre, M., Bosman, J., Roenneberg, T., Merrow, M., 2010, A circadian clock in *Saccharomyces cerevisiae*, PNAS 107 (5), 2043–2047.

Gangl, H., Tiefenbrunner, W., Pfliegler, W.P., Sipizki, M., Leitner, G., Tscheik, G., Lopandic, K., 2017, Influence of artificial intersecies yeast hybrids and their F1 offspring an the aroma profile of wine, Mitteilungen Klosterneuburg, 67. 68–83.

Hartung, J., Elpelt, B., 1999, Multivariate Statistik, Oldenburg Wissenschaftsverlag.

Hofbauer, J., Sigmund, K., 1984, Evolutionstheorie und dynamische Systeme. Mathematische Aspekte der Selektion. Parey, Berlin-Hamburg.

Jaenisch, R., Bird, A., 2003, Epigenetic regulation of gene expression: how the genome integrates intrinsic and environmental signals, Nature genetics 33, 245–254, doi:10.1038/ng1089..

Libkind D., Hittinger C.T., Valério E., Gonçalves C., Dover J., Johnston M., Gonçalves P., Sampaio J.P., 2011, Microbe domestication and the identification of the wild genetic stock of lager-brewing yeast, Proc. Natl. Acad. Sci. 108:14539–14544.

Lopandic K., Gangl H., Wallner E., Tscheik G., Leitner G., Amparo Querol A., Borth N., Breitenbach M., Prillinger H., Tiefenbrunner W., 2007, Genetically different wine yeasts isolated from Austrian vine-growing regions influence differently wine aroma and contain putative hybrids between *Saccharomyces cerevisiae* and *S. kudriavzevii*, FEMS Yeast Research 7: 953–965.

Lopandic K., Pfliegler W.P., Tiefenbrunner W., Gangl H., Sipiczki M., Sterflinger K., 2016, Genotypic and phenotypic evolution of yeast interspecies hybrids during high-sugar fermentation. Appl. Microbiol. Biotechnol. 100 (14), 6331–43. doi: 10.1007/s00253-016-7481-0. Epub 2016 Apr 13.

Nguyen H.V., Legras J.L., Neuveglise C., Gaillardin C., 2011, Deciphering the hybridization history leading to the Lager lineage based on the mosaic genomes of *Saccharomyces bayanus* strains NBRC1948 and CBS380, PLoS One 6:e25821.

Ninnemann, H., Butler, W.L., Epel, B.L., 1970, Inhibition of respiration in yeast by light, Biochem. Biophys. Acta 205, 499–506.

Pérez-Torrado R., González S.S., Combina M., Barrio E., Querol A., 2015, Molecular and enological characterization of a natural *Saccharomyces uvarum* and *Saccharomyces cerevisiae* hybrid, Int. J. Food Microbiol. 204, 101–110.

Pfliegler W.P., Atanasova L., Karanyicz E., Sipiczki M., Bond U., S. Druzhinina I.S., Sterflinger K., Lopandic K., 2014, Generation of New Genotypic and Phenotypic Features in Artificial and Natural Yeast Hybrids, Food Technology and Biotechnology 52, 46–57.

Robertson J.B., Davis, C.R., Johnson, C.H., 2013, Visible light alters yeast metabolic rhythms by inhibiting respiration, PNAS 110 (52), 21130–21135.

Roenneberg, T., Merrow, M., 2001, Seasonality and Photoperiodism in Fungi, Journal of Biological Rhythms 16 (4), 403–414.

Russo, V. E., 1988, Blue light induces circadian rythms in the bd mutant of *Neurospora*: Double mutants bd,wc-1 and bd,wc-2 are blind. Photochem. Photobiol 2, 59–65.

Stübiger G., Wuczkowski M., Mancera L., Lopandic K., Sterflinger K., Belgacem O., 2016, Characterization of Yeasts and Filamentous Fungi using MALDI Lipid Phenotyping, Journal of Microbiological Methods 130, 27–37.

Sulkowski, E., Guerin, B., Jaques, D., Slonimski, P.P., 1964, Inhibition of protein synthesis in yeast by low light-intensities of visible light, Nature 202, 36–39.

Woodward J.R., Cirillo, V.P., Edmunds Jr., L.N., 1978, Light Effects in Yeast: Inhibition by Visible Light of Growth and Transport in *Saccharomyces cerevisiae* Grown at Low Temperatures, J. Bacteriol. 133 (2), 692–698.

